# A metagenomic view on fungal diversity in freshwater lakes

**DOI:** 10.1101/2022.06.20.496890

**Authors:** Anushree Sanyal, Mariana Kluge, Miguel Angel Redondo, Moritz Buck, Maliheh Mehrshad, Sarahi L Garcia, Stefan Bertilsson, Sari Peura

## Abstract

Fungi are essential components in a wide range of ecosystems and while major efforts have been spent on disentangling the diversity and functional roles of fungi in terrestrial environments, our knowledge about aquatic fungi is lagging. To address this knowledge gap, we explored metagenomes from 25 lakes from the arctic and boreal zone and one tropical reservoir with the aim of describing the community structure of fungi and fungi-like organisms (Oomycota). A second objective was to identify possible environmental factors influencing the composition of the fungal communities. Our results show that the main fungal phyla and orders are the same across all the lakes despite the differences in geographic location and prevailing climate. Still, there was differential distribution of more highly resolved taxa across the lakes that accordingly featured distinct communities, possibly caused by differential availability of carbon substrates in the lakes. A more detailed classification of sequences related to the pathogenic Oomycota genus *Phytophthora* clearly demonstrated that while technologies now exist for sequencing entire microbial communities in great detail, we are still severely limited by insufficient coverage of eukaryotic sequences and genomes in public databases.

## Introduction

Fungi are found in almost all habitats on Earth, from the human body to sediments in Antarctica (Huffnagle and Noverr 2013, Selbmann et al. 2015) and approximately 2% of all biomass on Earth has been estimated to be of fungi (Bar-On et al. 2018). In terrestrial environments, it has for quite some time been established that fungi mediate central and indispensible processes in nutrient cycling and hold key roles as decomposers, saprophytes, parasites, symbionts and necrotrophs (Webster and Weber 2007, Gargas et al. 1995, Taylor et al. 2015, Peay et al. 2016). However, for aquatic ecosystems, both the diversity of fungi and their ecological and biogeochemical roles are much less understood (Grossart and Rojas-Jimenez 2016, Grossart et al. 2019). Early DNA-based studies on marine microeukaryotes found that fungi comprised only about 1% of the combined marine eukaryote communities (Massana and Pedros-Alio 2008), and because of this they have typically been overlooked in studies of marine microeukaryotes and considered as being ecologically and biogeochemically insignificant. However, recent findings have demonstrated that the apparent low abundance of aquatic fungi in molecular microeukaryote datasets and surveys was due to a general lack of representative entries in the commonly used databases (Khomich et al. 2018) and that fungi can comprise up to 33 % of aquatic microeukaryote communities (Lèpere et al. 2016). More extensive use of high-throughput sequencing over the last decade has revealed that fungal diversity in aquatic systems has been severely underestimated (Scheffers et al. 2012), and that a substantial but variable proportion of eukaryotic sequences are in fact fungal (Richards et al. 2012, De Vargas et al. 2015, Deboras et al. 2017). In agreement with this, many recent studies have found that fungi are abundant and diverse in both marine (Lèpere et al. 2015, Gladfelter et al. 2019) and freshwater (Ishii et al. 2015, Duarte et al. 2015, Lèpere et al. 2016) microbial communities, with their mere quantitative significance implying that they also play significant ecological and biogeochemical roles in these habitats (Wurzbacher et al. 2018). Nevertheless, data on the composition of fungal communities are still scarce for inland lakes (Grossart et al. 2019).

Aquatic fungi are, by definition, fungi that rely on aquatic habitats for either their entire life cycle or parts thereof. They are categorized into three broader groups based on their reliance on aquatic habitats, extent of adaptation and activity: indwellers, periodic immigrants and versatile immigrants. Indwellers are completely adapted and continuously active aquatic fungi and their whole life cycle of conidium production, release and dispersal typically occurs under water (El-Elimat, et al. 2021). The periodic immigrants are less adapted to aqueous habitats and occur across both terrestrial and aquatic environments and are periodically active. The versatile immigrants are the least adapted to water and only occasionally active, while the transients are those that exhibit no activity in natural waters (Park 1972, Grossart et al. 2019). Classification has also been done based on functional traits (Wurzbacher et al. 2010, Krauss et al. 2011, Grossart et al. 2021), which might be more appropriate than phylogeny to understand the ecological and biogeochemical role of fungi in aquatic systems. These aquatic fungal groups include aquatic hyphomycetes, chytridiomycota and yeasts. Aquatic hyphomycetes are a heterogeneous group typically found in plant litter of streams (Ingold, 1942; Koivusaari et al. 2019). Chytridiomycota are parasitic and saprophytic fungi typically found in pelagic zones of stagnant waters (Wurzbacher et al. 2010), Yeasts are ubiquitous in freshwater ecosystems, especially in the pelagic zone of lakes (Wurzbacher et al. 2010). While not fungal in the strict sense, fungal-like oomycetes are mainly saprobes, and some of them are important animal parasites (e.g., on fish and crustaceans) or plant pathogens (Shearer et al. 2007).

While most studies that have informed us about the ecology and diversity of aquatic fungal communities have been based on high throughput sequencing of either amplified 18S ribosomal RNA genes (Lepère et al. 2019) or the intergenic ribosomal RNA spacer (ITS; Monard et al. 2016), many of the unknown fungi might not amplify with existing biomarker primers and will thus go unnoticed (Kounosu et al. 2019). To overcome this limitation, we explored the use of shotgun metagenome sequencing for characterizing fungal and oomycete community composition. To this end we included samples from 25 lakes distributed across the boreal and arctic region and one tropical reservoir. Our aims were to investigate whether environmental conditions are related to the fungal community composition and which are the main phyla and orders found in these lakes. Additionally, we scrutinized the presence and diversity of genus *Phytophthora* for a more detailed analysis on a species level. This is a group of fungal-like organisms that comprises important pathogens of crops and trees (Erwin and Ribeiro, 1996) and recent studies exist on the species distribution of this genus in water bodies in northern latitudes (Redondo et al. 2018a,b).

## Materials and Methods

### Sample and data processing and chemical analyses

The sample collection and metagenome sequencing of the lake samples used in this study are thoroughly described in Buck et al. (2021). In short, pelagic samples were collected from 26 freshwater lakes (Table 1) ranging in geographic location from the arctic to the tropics (hereafter referred to as “lake dataset”). Microbial cells were collected by filtration onto a 0.2 µm filter and for most of the waterbodies samples from multiple depths were included, representing a gradient of decreasing oxygen and increasing nutrient concentration from surface to bottom. Samples for analysis of relevant chemical variables were collected simultaneously. For one of the lakes (Lake Mekkojärvi), a multi-year time series for the ice free seasons 2011-2013 was included to assess the stability of the fungal community (hereafter referred to as “time series data”) and for two other lakes (Alinen Mustajärvi and Lomtjärnan) samples from multiple years and multiple depths were included to estimate the stability of vertical distribution of different fungal taxa (hereafter referred to as “vertical stability data”). For all samples, the total DNA was extracted using PowerSoil DNA extraction kit according to the manufacturer’s instructions. After quality control and library preparation, the samples were sequenced on an Illumina NovaSeq instrument hosted by the Science for Life Laboratory National Genomics Infrastructure at Uppsala University, Sweden.

**Table 1.** The mean, maximum and minimum proportions of unclassified, fungal and Oomycota reads detected in each of the lakes with multiple samples and for the lakes with one sample only, the absolute proportions.

| Lake | Unclassified reads |  |  | Fungal reads |  |  | Oomycota reads |  |  |
| --- | --- | --- | --- | --- | --- | --- | --- | --- | --- |
|  | Mean (%) (SD) | Max (%) | Min (%) | Mean (%) (SD) | Max (%) | Min (%) | Mean (%) (SD) | Max (%) | Min (%) |
| Alinen Mustajärvi | 52.6 (7.7) | 65.4 | 38.3 | 3.7 (2.2) | 6.2 | 1.8 | 0.12 (0.11) | 0.28 | 0.03 |
| Björntjärnan | 44.2 (5.6) | 49.9 | 38.8 | 2.4 (0.8) | 3.3 | 1.7 | 0.09 (0.06) | 0.14 | 0.03 |
| Erken | 46.5 (12.9) | 55.6 | 33.8 | 1.9 (0.9) | 4.6 | 1.3 | 0.07 (0.06) | 0.17 | 0.03 |
| Haukijärvi | 44.3 (8.1) | 56 | 38 | 2.7 (1.4) | 4.8 | 1.9 | 0.13 (0.15) | 0.34 | 0.03 |
| Keskinen Rajajärvi | 56.1 (8.2) | 68.5 | 49.3 | 3.6 (1.4) | 5.9 | 2.5 | 0.11 (0.07) | 0.21 | 0.04 |
| Ki1 | 49.3 (11.8) | 63.1 | 35.7 | 3.1 (1.5) | 4.5 | 1.4 | 0.16 (0.13) | 0.3 | 0.02 |
| Ki2 | 41.8 (4.2) | 45.3 | 36.5 | 2.1 (0.4) | 2.3 | 1.5 | 0.07 (0.03) | 0.1 | 0.03 |
| LaPlata | 43.8 (4.1) | 48.4 | 37.4 | 2.5 (0.5) | 3.2 | 1.8 | 0.05 (0.02) | 0.07 | 0.03 |
| Lomtjärnan | 43.8 (5.1) | 50.8 | 35.8 | 2.6 (1.6) | 3.8 | 1.5 | 0.1 (0.11) | 0.17 | 0.03 |
| Lovojärvi | 44.3 (8.6) | 63.1 | 37.8 | 2.7 (1.2) | 5.4 | 1.9 | 0.07 (0.06) | 0.2 | 0.03 |
| Mekkojärvi | 37 (3.7) | 52.3 | 30.7 | 1.5 (0.8) | 2.8 | 1.1 | 0.03 (0.04) | 0.09 | 0.01 |
| Närtjärnen | 46 (6.4) | 52.9 | 37.5 | 3.1 (1.2) | 4.5 | 1.7 | 0.15 (0.14) | 0.36 | 0.03 |
| Odrolstjärnen | 40 (1.8) | 41.2 | 38.7 | 2 (0.2) | 2.2 | 1.8 | 0.1 (0.03) | 0.12 | 0.08 |
| Umea3 | 44.7 (3.3) | 48.5 | 40.2 | 2.3 (0.4) | 2.8 | 1.8 | 0.07 (0.04) | 0.12 | 0.03 |
| Vähä-Keltajärvi | 57.8 (12.8) | 70.9 | 46.8 | 4 (2) | 6 | 2.2 | 0.1 (0.06) | 0.19 | 0.05 |
| Ylinen Rajajärvi | 52.1 (14.5) | 64.3 | 35.7 | 3.2 (1.8) | 4.7 | 1.3 | 0.11 (0.09) | 0.22 | 0.02 |
| Lakes with one surface sample | Unclassified reads (%) |  |  | Fungi (%) |  |  | Oomycetes (%) |  |  |
| Bengtsgölen | 43.9 |  |  | 2.2 |  |  | 0.07 |  |  |
| Fyrsjön | 61.8 |  |  | 4.7 |  |  | 0.18 |  |  |
| Glimmingen | 56.2 |  |  | 3.9 |  |  | 0.2 |  |  |
| Långsjön | 39.4 |  |  | 2.4 |  |  | 0.08 |  |  |
| Lillsjön | 68.4 |  |  | 7.8 |  |  | 0.38 |  |  |
| Lotsjön | 63.4 |  |  | 4 |  |  | 0.16 |  |  |
| Malstasjon | 55.2 |  |  | 3.6 |  |  | 0.12 |  |  |
| Parsen | 60.2 |  |  | 4 |  |  | 0.11 |  |  |
| Plåten | 56 |  |  | 3.3 |  |  | 0.12 |  |  |
| Stortöveln | 50.8 |  |  | 3.8 |  |  | 0.18 |  |  |

For each of the samples, a set of relevant descriptive environmental variables were also measured. These included dissolved oxygen (DO), total organic carbon (TOC) and total nitrogen (TN). The analyses have been thoroughly described in Buck et al. (2021).

### Analysis of fungal reads

Combined, our metagenome dataset consisted of 1.28 TB of data (for more details, see Buck et al. 2021). The raw data were first trimmed using Trimmomatic, after which all the reads were taxonomically classified using Kaiju with an inhouse database including the NCBI nr database combined with the Joint Genome Institute MycoCosm database (Grigoriev et al. 2012). Kaiju was run in greedy mode where the thresholds of the minimum match score were set at 75 and 5 mismatches were allowed. A large proportion of the reads (31.1 to 76.7%) remained unclassified in the Kaiju output and those were excluded from further analyses (Table 1). In order to analyze the composition of the fungal and fungi-like communities in the lakes, we included in our analyses all reads that were classified as fungi or oomycota. For these we estimated the proportion of fungal and oomycetal reads relative to all the classified reads. Then, a table with counts of all reads classified as fungal and oomycotal orders was compiled. Prior to further analysis, the table was normalized to the number of reads in the least represented sample (162 786 reads) to account for the variable sequencing depth across the samples. For statistical tests involving alpha diversity, taxa with less than 1000 reads in the combined normalized dataset were discarded. For beta diversity and PERMANOVA analyses the data was Hellinger transformed.

The abundances of different *Phytophthora* species were evaluated by retrieving all the reads assigned to this genus and classifying them on the species level using Kaiju and the in-house database. These counts were normalized by calculating the proportions of each of the species for each of the samples.

### Statistical analysis

Alpha diversity indices of fungal and oomycetal communities were calculated from the normalized table using six different indices related to diversity: Shannon, Pielou’s and inverse Simpson indices, number of observed taxa, Chao1, and abundance-based coverage estimator (ACE). The difference in diversity between lakes was tested using Kruskal-Wallis test with Dunn’s test as post-hoc test (correction for multiple testing using Bonferroni method) for indices with significant difference between lakes and layers. Beta diversity, in terms of the variability in community composition among lakes, was visualized with a nonmetric multidimensional scaling (NMDS). Also, for each depth layer, beta-diversity was assessed with a permutational analysis of multivariate dispersions (PERMDISP, betadisper function of vegan package) to check the deviations of each sample from the group centroid. The differences between groups were verified by the permutest function in the vegan package. The adonis function of the vegan package (Oksanen et al. 2019) was used for PERMANOVA analysis to determine whether lake and depth layer significantly impacted the composition of the community, and pairwise.adonis function was used to check the differences between groups. The differences in community composition were visualized with an NMDS. For a subset of 52 samples for which TOC, TN and O_2_ data were available, PERMANOVA (marginal effects) and Mantel tests were applied to analyze the relationships between total community composition and these environmental factors. Correlation between oxygen concentration, which was available for all samples, and fungal abundance was tested using Spearman rank correlation.

## Results

### Diversity of fungal and fungal-like communities differs between freshwater lakes

The proportion of fungal reads in the classified sequences across the samples ranged from 1.1 % at the anoxic bottom layers of the lakes (hypolimnion; M = 2.3 %, SD =1.4) to 11.4 % in the oxygen-rich surface layer (epilimnion; M = 4.0 %, SD = 1.5). Of these reads, those assigned as originating from oomycetes comprised ≤ 0.2% in the hypolimnion and ≤ 0.4% in the epilimnion across all lakes (Table 1 = 1). Despite the seemingly decreasing trend in the fungal abundance from the lake surface to the bottom, with regards to average abundances, there was no significant correlation between fungal or oomycetal abundance and oxygen concentration. The overall fungal and fungal-like community composition at the order level was different between lakes (Fig 1a; PERMANOVA: *p* < 0.001, *F* = 2.9, R^2^ = 0.34), and layers (Fig 1b; *p* < 0.001, *F* = 13.7, R^2^ = 0.12). Pairwise analyses between depth layers showed that the hypolimnion communities were significantly different (*p* < 0.05) from the epiand metalimnion. There was a slight decreasing trend in alpha diversity from the surface to the bottom layer, but the difference was not significant (Supplementary Fig S1a). There was no significant difference in the community composition, i.e. beta diversity, between the depth layers, as shown by the analysis of dispersion. The following alpha diversity indices were significantly different between lakes and layers: Shannon (lakes: χ^2^ = 55.1, *p* = 0.001; layers: χ^2^ = 20, *p* = 0.0000449), inverse Simpson (lakes: χ^2^ = 54.3, *p* = 0.001389; layers: χ^2^ = 20.2, *p* = 0.00004105) and Pielou’s (lakes: χ^2^ = 55.6, *p* = 0.0009781; layers: χ^2^ = 19.5, *p* = 0.00005745). Posthoc tests for Shannon, inverse Simpson and Pielou’s were not significant between lakes after bonferroni corrections. Posthoc tests showed that hypolimnion significantly differed (*p* < 0.05) from other layers for Shannon, inverse Simpson and Pielou’s.

**Figure 1.**
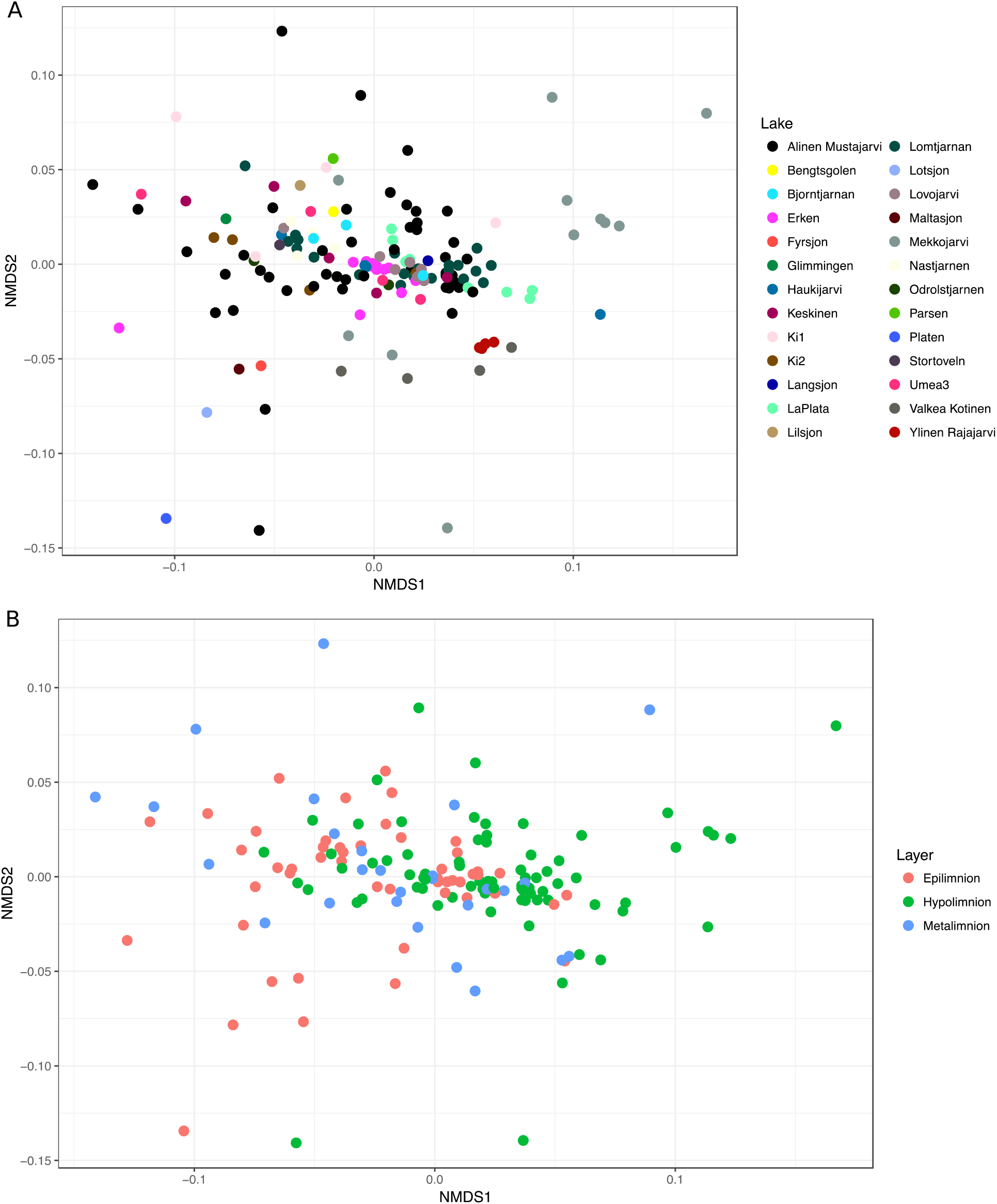
NMDS illustrating the relative abundance of the taxonomic composition of the fungal communities across the lakes, by A) lake and B) layer. Stress = 0.16.

We also investigated which environmental factors (TOC, TN and O_2_) best explained the community composition at the order level in a subset of 52 samples in the lake dataset. TOC (*p* = 0.000999, *F* = 5.01) and TN (*p* = 0.0003996, *F* = 3.92) explained 8 and 6.3 % of the variation, respectively. O_2_ was also significant (*p* = 0.016983, *F* = 2.65) and explained 4.3 % of the variation. Subsequent Mantel tests showed that only TOC was significantly correlated (*p* = 0.0003) with the community composition, explaining 26 % of the variation. When considering only the oomycete communities, only O_2_ was significantly associated with community composition (*p* = 0.024, *F* =4.18; R^2^ = 0.073).

### Community composition

In terms of fungal phyla, the most abundant phylum across all lakes was Ascomycota, which contributed up to 67.9 % of all fungal reads in the individual samples (Fig 2, Table 1 = 1; M = 47 %, SD = 4.7). This was followed by Basidiomycota (up to 44.2 %, M = 37.1 %, SD = 4.0) and Chytridiomycota which contributed less than 10 % to the individual samples (M = 4.7 %, SD = 1.5), with the exception of one sample with 15.5 % of such reads. Also, Mucoromycota, Zoopagomycota, and Blastocladiomycota were found in most lakes, with mean abundances of 5.3 % (SD =1.2), 2.5 % (SD = 0.7) and 0.6 % (SD = 0.2), respectively. Interestingly, the tropical reservoir (La Plata) shared a similar overall community composition as the boreal lakes (Fig 2 and Fig 3). Finally, the mean proportion of reads belonging to the oomycota across all lake samples was 0.09 % (SD = 0.09).

**Figure 2.**
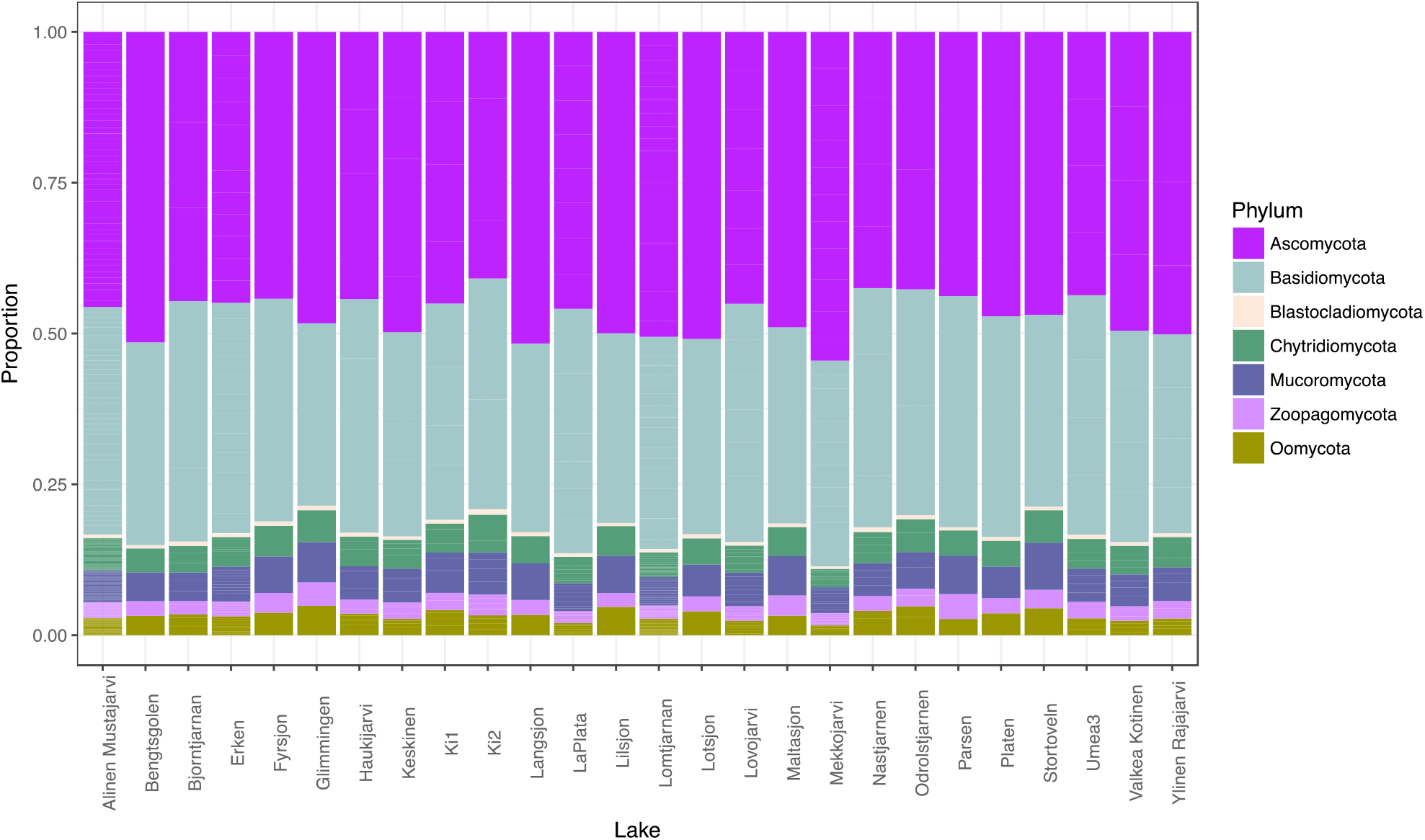
The proportions of the fungal phyla and Oomycota across the lakes. Most of the lakes had samples taken from multiple depths and for these, the proportions are averages across all the samples from the same lake. The thin lines within each phylum represent the contribution of each of the samples to the average proportion of that phylum, thus, the lakes with no lines only had one sample available.

**Figure 3.**
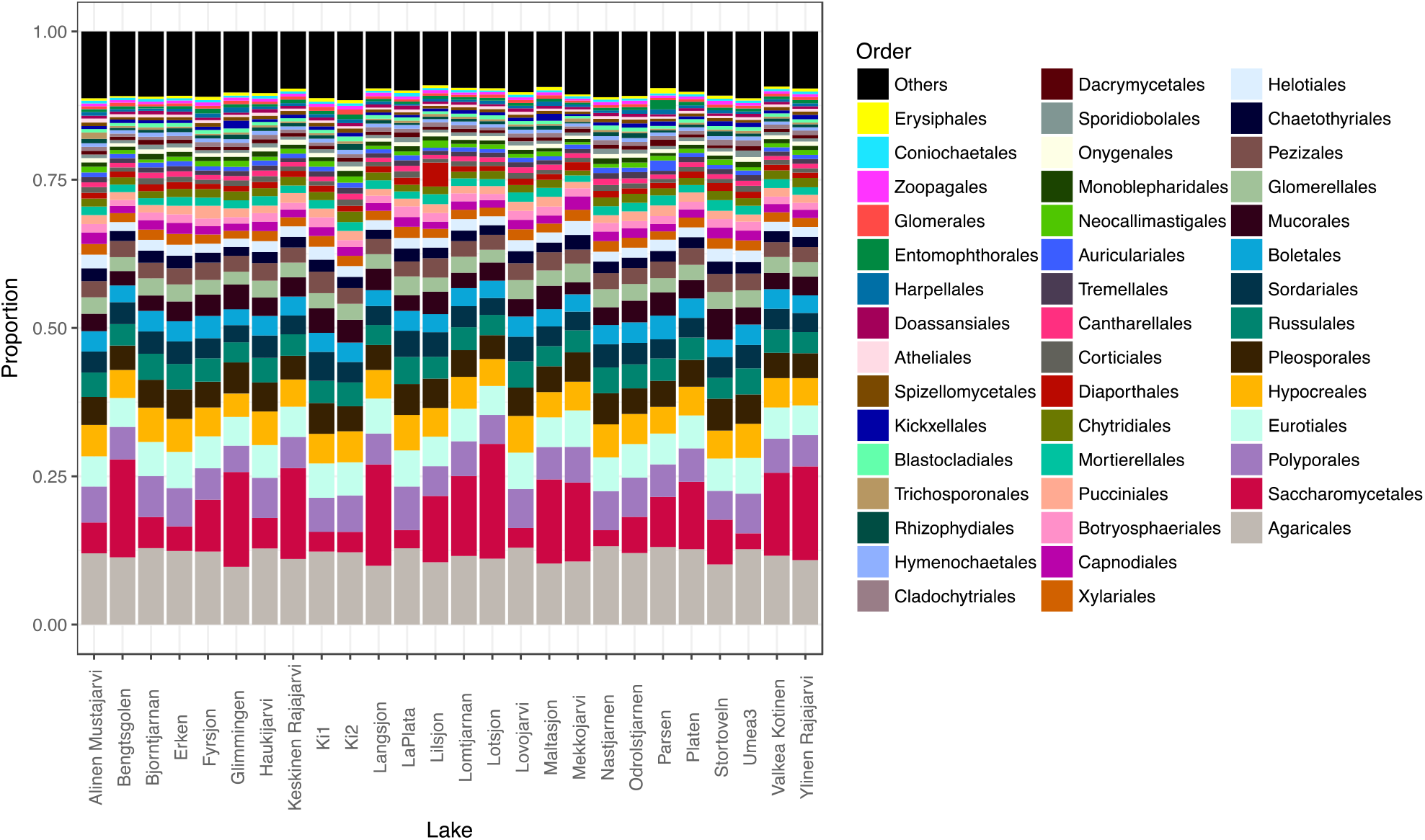
The proportions of all orders with more than 100 000 reads in the lakes data. Most of the lakes had samples taken from multiple depths and for these, the proportions are averages across all the samples from the same lake.

Regarding the fungal orders, the 45 fungal and two oomycotal orders represented by more than 100 000 reads in the combined lake metagenome are shown in Fig 3. The number of orders found in individual lakes ranged from 28 to 56 when orders represented by at least 1000 reads were included. Of these orders, 25 were present in all lakes (Table 2).

**Table 2:** The most abundant fungal and oomycotal orders represented by more than 100000 reads

| Order | Number of reads |
| --- | --- |
| Agaricales* | 2935915 |
| Saccharomycetales* | 1913521 |
| Polyporales* | 1500442 |
| Eurotiales* | 1351694 |
| Hypocreales* | 1304604 |
| Pleosporales* | 1158517 |
| Russulales* | 993432 |
| Sordariales* | 896178 |
| Boletales* | 801329 |
| Mucorales* | 724435 |
| Glomerellales* | 688621 |
| Pezizales* | 663614 |
| Chaetothyriales* | 498389 |
| Helotiales* | 494340 |
| Xylariales* | 425287 |
| Capnodiales* | 420912 |
| Botryosphaeriales* | 344929 |
| Pucciniales* | 342841 |
| Mortierellales* | 339228 |
| Peronosporales* | 319885 |
| Chytridiales* | 314647 |
| Saprolegniales* | 292887 |
| Diaporthales* | 261578 |
| Corticiales | 220080 |
| Cantharellales* | 207146 |
| Tremellales* | 205209 |
| Auriculariales | 193448 |
| Neocallimastigales | 186636 |
| Monoblepharidales | 181429 |
| Onygenales | 171170 |
| Sporidiobolales | 159085 |
| Dacrymycetales | 157338 |
| Cladochytriales | 155185 |
| Hymenochaetales | 150438 |
| Rhizophydiales | 144380 |
| Trichosporonales | 142562 |
| Blastocladales | 138067 |
| Kickxellales | 132425 |
| Spizellomycetales | 132404 |
| Atheliales | 123525 |
| Doassansiales | 122627 |
| Harpellales | 121299 |
| Entomophthorales | 118546 |
| Glomerales | 115580 |
| Zoopagales | 112155 |
| Coniochaetales | 105536 |
| Erysiphales | 103069 |

\*indicates the orders present in all lakes.

Overall, each order had very similar proportions across all the lakes, with rather minor variations. The most variable order was Saccharomycetales, which was the second most abundant order across the lakes. The most abundant order was Agaricales followed by Saccaromycetales, Polyporales, Eurotiales, and Hypocreales (Fig 2). There were also 16 other orders that were consistently present in all lakes, with abundances between 1% and 5%. In addition, four orders of the Oomycota (Peronosporales, Saprogeniales, Albuginales and Pythiales) were found consistently in all 26 lakes.

### Phytophthora

As Oomycetes includes prominent plant pathogens, some of which are known to be dispersed from one area to another via waterways, we looked more closely into the diversity and distribution of one important group of oomycete pathogens, namely *Phytophthora*. Among the reads classifies as *Phytophthora*, the most abundant species identified by Kaiju were *P. megakarya* (M = 11.4 %, SD = 4.6), *P. parasitica* (M = 9.4 %, SD = 5.0), *P. sojae* (M = 8.0 %, SD = 2.0) and *P. cactorum* (M = 7.1 %, SD = 4.2).

### Stability of the fungal communities over time

Samples from three of the lakes were collected for multiple years, including a more intensive sampling from hypolimnion at Lake Mekkojärvi (Fig S2a; time series data) and repeated samplings from lakes Lomtjärnan, and Alinen Mustajärvi (Fig 4; vertical stability data). In all the lakes with time series data, the most abundant taxa were consistent with the observations from the complete lake dataset. The abundance of fungi in the hypolimnion of Lake Mekkojärvi was low but rather stable over the three years, with average proportions of fungal and oomycetes reads from all classified reads in the time series being 0.9 % (SD = 0.1) and 0.01 % (SD = 0.003), respectively (Fig S2b). This extensive sampling effort at Mekkojärvi showed that in general the composition of fungal community is rather stable, with the same taxa present throughout the different time periods. It also showed that the community composition was similar to the the lake dataset, and the main source of variation was the proportion of Saccharomycetales which was greater in some samples. The time series included duplicate samples for most time points, but in most cases the increased abundance of Saccharomycetales was observed in only one of the replicates, suggesting that their distribution is patchy and possibly associated with particles.

**Figure 4.**
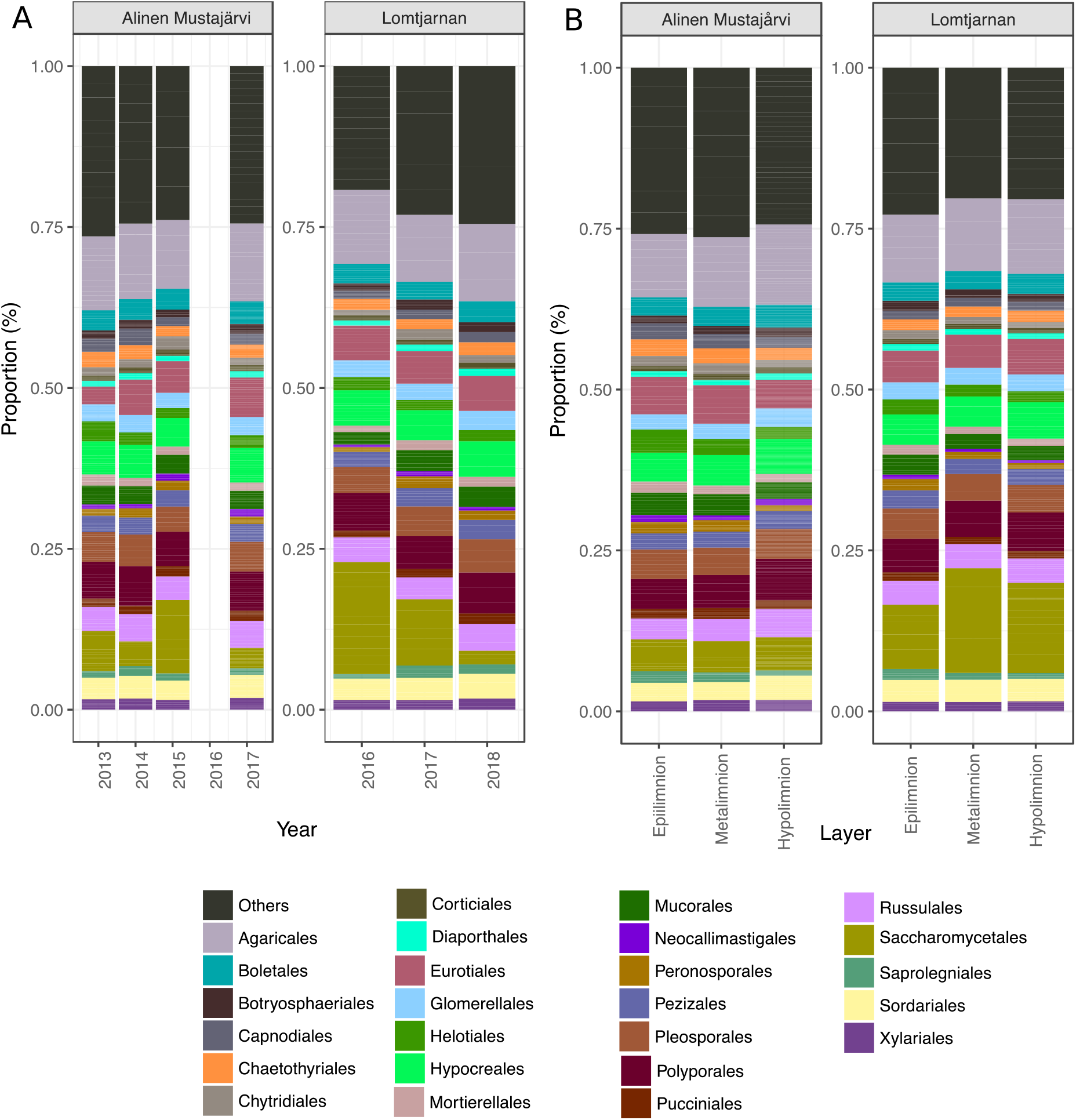
The composition of the fungal communities in the order level in the yearly samples from lakes Alinen Mustajärvi and Lomtjärnan by A) year and B) layer

Also, the community composition in lakes Alinen Mustajärvi and Lomtjärnan that had been sampled on multiple occasions during different years, had the same high variation for the abundance of Saccharomycetales (Fig 4), while other orders were less variable. However, compared to interannual variation, the composition of different layers of these lakes did not have a clear pattern, but the relative abundance of Saccharomycetales was rather stable across the depth profiles (Fig 4).

## Discussion

Overall, our results suggest that the fungi comprise a small but stable part of microbial communities in boreal lakes. The low contribution of fungi to the total microbial community is in line with a recent study reporting on the distributions of bacteria, archaea and eukaryotes in thermokarst ponds (Wurzbacher et al. 2017). Studies are typically focused on only one type of microbes (bacteria, archaea, algae or fungi) at a time. This approach is justifiable due to methodological difficulties in simultaneously addressing such a genetically diverse community, but it leaves open the question on the relative abundances of different microbial organisms. To our knowledge one study has attempted to show the contribution of fungi to the total microbial community along a terrestrial-freshwater gradient where they found habitat generalist and habitat specific species (Monard et al. 2016). This study found that the fungi of the orders Capnodiales and Helotiales were most abundant in freshwaters and mineral soils. These orders were also among the most abundant in our lake dataset (Table 2). In our study, which, to our knowledge, is the first study using shotgun sequencing to study fungal diversity specifically in freshwaters, we could get estimates on the abundance of fungi in relation to other members of the microbial community and it showed that the fungi comprise a low but stable proportion of the communities seen in all of the analyzed samples from different lakes, time series and vertical profiles. Based on our data, we cannot deduce whether these fungi are active, but a previous study on lake fungi using 18S rRNA transcripts has suggested that 13 % of fungal OTUs would be active (Lepére et al. 2019). Considering that the other members of the lake microbial communities present similar activity patterns (e.g. for freshwater bacteria estimates range from 6 to 33 % active (Chrzanowski et al. 1984, del Giorgio and Scarborough 1995)), and since some of the samples did have fairly high fungal contribution, with up to 11 % of the reads in the oxic layer of Lake Alinen Mustajärvi, suggest that fungi can contribute to the total metabolism in the lake.

Both the lake and depth layer had a significant effect on the community composition at the order level with the communities residing in hypolimnetic waters being distinct from the other layers, even though a few orders were dominant throughout the water column. Oxygen was still not a main driver of community composition, suggesting that oxygen depletion in the hypolimnion may not be a structuring factor underlying the shifting community structure with increasing depth. Our results suggest that among all the variables included in the analysis, organic substrates (with TOC as proxy) was the most influential factor explaining the variable community composition. However, admittedly our set of environmental variables was limited and there may well be other factors that are more important than organic carbon. Furthermore, our data does not allow to infer about the effects of the quality of carbon. In agreement with our findings, a study that included 77 oligotrophic lakes in Scandinavia also found significant correlation between TOC and aquatic fungal community composition (Khomich et al. 2017), and this pattern was also corroborated in a recent study on thermokarst ponds revealing a significant impact of both carbon quality and quantity on fungal community composition (Kluge et al. 2021). Considering that fungi generally are expected to be efficient degraders of various carbon compounds (Phillips et al. 2013, Frey 2019), it is not surprising that organic substrate availability would have an impact on fungal community composition.

Our results show that the community composition varied between layers, with significant differences in the alpha diversity indices across the layers and the same fungal orders were dominant in all lakes. Ascomycota was the dominant phylum in all of the lakes, followed by Basidiomycota, Chytridiomycota and Mucoromycota. These results are in agreement with other studies on lakes that have found the Ascomycota to be the dominant phylum (Lepére et al., 2019, de Souza et al. 2021). However, in a study in oligotrophic lakes in Scandinavia, the Chytridiomycota represented the most abundant species, whereas Ascomycota and Basidiomycota represented the largest proportion of detected species (Khomich et al. 2017). Chytridiomycota, along with Cryptomycota, were also found to dominate fungal communities in ice-covered lakes in Antarctica (Rojas-Jimenez et al. 2017). Rather than representing true biogeographical patterns, such divergent findings may rather reflect the poor coverage of aquatic fungi in current databases, making it difficult if not impossible to know how much of the fungal community diversity is overlooked. Over the past years, many newly described ascomycetes were reported in freshwaters (El-Elimat et al. 2021), revealing that there is still much to be revealed about their diversity. A similar scenario is seen in the case of the poorly known lineages of Chytridiomycota and Rozellomycota, which are grossly underrepresented in the current databases (Grossart et al. 2019, Frenken et al. 2017). Another explanation for apparent contrasting patterns in fungal biodiversity is the use of different primers across studies. All of the previous studies cited here are based on amplicons and different primer sets may be biased towards or against specific fungal groups (Tedersoo and Lindahl, 2016).

Ascomycota, the dominant phylum in our study, includes > 3000 species that are specialized on a saprobic lifestyle in freshwater habitats where they successfully grow and sporulate (Shearer et al. 2007, Kirk et al. 2008, Shearer and Raja 2010). In line with this, many of the most dominant orders in our study belonging to the phylum Ascomycota were those known to represent aquatic fungi, such as Eurotiales, Hypocreales, Pezizales, Diaporthales, Sordariales, Xylariales, Mucorales, Chaetothyriales, Helotiales, Saprolegniales, Peronosporales and Pleosporales (Shearer et al. 2007, Shearer and Raja 2010, Embree and Indoh 1967, Chandrasiri et al. 2021, Kohout et al. 2012, Beakes and Sekimoto 2009), and could represent indwellers in the lakes. Aquatic Saprolegniales are widespread animal pathogens of salmonids, amphibians and crustaceans (Bruno et al. 2011, Hussein and Hatai, 2002, Fregeneda-Grandes et al. 2007). Based on the ubiquity of the taxa across the study lakes, we speculate that Chytridiales, Rhizophydiales, Blastocladiales, Neocallimastigales, and Cladochytriales could also be indewellers, in line with previous studies (Lepère et al. 2008, Kagami et al. 2014, Lefèvre et al. 2012, Mozley-Standridge et al. 2009, Steiger et al. 2011, Money 2016). Members of Glomerellales have been identified in seagrass tissues and could be speculated to be indewellers (Ettinger and Eisen 2019). In contrast, Agaricales, Polyporales, Boletales, Russulales, Mortierellales and Botryosphaeriales within the phylum Basidiomycota which were among the most abundant orders found in our lakes, is more likely to be an inactive community member. While there are some true aquatic Agaricales and Mortierellales (Frank et al. 2010, Embree and Indoh 1967), this order typically comprises terrestrial fungi. Manymembers of Russulales, Cantharellales and Boletales are mycorrhizal fungi which depend on plant hosts to sustain growth (Halling 2001, Bonito et al. 2013, Xu et al. 2021, Floudas 2021). Most members of the Polyporales are wood composers (Money 2016), members of Mortierellales, Tremellales and Cantharellales are saprobes in soil, and members of Botryosphaeriales, Tremellales and Pucciniales are known as plant pathogens (Batista et al. 2021, Helfer 2014, Gams et al. 2004, van der Klei et al. 2011). When such mycorrhizal fungi, wood decomposers, soil saprobes and terrestrial plant pathogens are detected in aquatic habitats, they probably enter the aquatic system through run off from the terrestrial ecosystem (Khan 1993). Members of Capnodiales have been found associated with roots of submerged aquatic plants; studies have suggested their adaptation to halophilic conditions (Kohout et al. 2012, Gunde-Cimerman et al. 2000). Thus, we speculate that part of the Agaricales, Polyporales, Boletales, and Russulales found in the lakes likely represent transient taxa, which would arrive and decline overtime and likely do not have any major ecological role in aquatic systems except as sources of nutrients (Park 1972, Kohout et al. 2012). This is also a consideration that needs to be taken into account for the whole dataset; when a survey is based on DNA extracted from the water, a variable portion of this material may originate from remnant DNA from organisms that are no longer alive. Thus, there is a possibility, that in general many of the detected taxa are due to fungal DNA from the watershed rather than actively replicating communities residing in the lakes. However, if a major proportion of the taxa would be coming from the surrounding soil and be inactive in these lakes, we would expect a larger variation in the community composition across the climatic zones, as it has been shown that the fungal community composition at the order level varies significantly between arctic, boreal and tropical soils (Tedersoo et al. 2014). This suggests that the community composition in our lakes is not to any larger extent shaped by fungal imports from the surrounding watershed.

Interestingly, the proportion of the Saccharomycetales order was particularly variable across the lakes. These are saprotrophic ascomycete yeasts (Suh et al. 2006) known for their ability to ferment sugars (Kurtzman 2011). In the time series dataset where we had duplicate samples from the same time point but from different locations in the lake, most time points with increased abundance of Saccharomycetales had higher proportion for only one of the two samples. This suggests that the changes in the relative abundance of this taxon could be very local and could be related, for example, to particles carrying a large number of spores or organic particles that undergo active fungal degradation rather than a systematic increase in the relative abundance across the entire lake or water mass. While our data does not allow us to resolve the factors behind the variation in the relative abundance of Saccharomycetales, it does suggest that this is a dynamic taxon that could be important for total lake metabolism and should thus be scrutinized further

Many of the fungal orders found in the lakes contain well-known pathogens to various organismal groups which have obvious direct and indirect effects on the environment (Fisher et al. 2012, King and Lively 2012). For example, members of orders Hypocreales, Chytridales and Rhizophydiales are known to be pathogens and parasites of insects, reptiles and amphibians (Wellehan and Divers 2019, Razzaghi-Abyaneh et al. 2015, Lefèvre 2007, 2012), while members of Catenariaceae of the order Blastocladiales are known to be pathogens of aquatic animals (Rand 2004) and members of Pichiaceae of the order Saccharomycetales are pathogenic to fish. A noteworthy group that was relatively abundant in our data was *Phytophthora* of the fungi-like oomycota. The genus *Phytophthora* contains several aggressive plant pathogens which disperse through water bodies, threatening the surrounding forest ecosystems (Bjelke et al. 2016, Hansen et al. 2000, Redondo et al. 2018a). In this study, we classified this genus down to species level to see if the classification corresponds to the expected composition of the *Phytophthora* community. Some of the identified species such as *P. cactorum* and *P. kernoviae* are pathogens of forest and woodland species. However, we found that 9 out of the 12 identified species are crop pathogens, even though the lakes were not in the vicinities of crop fields. Unexpectedly, none of the identified species belonged to the clades 6 or 9, which have a predominantly saprotrophic lifestyle in water bodies worldwide (Aram and Rizzo 2018, Hansen et al. 2012, Marano et al. 2016) and whose diversity is associated with water parameters such as pH and total nitrogen (Redondo et al. 2018b). These results suggest, that while the fungal community composition at the phylum and order levels match the expected taxa, inventories extended all the way to the species level may generate spurious results, as suggested by the *Phytophthora* species annotation. A closer scrutiny of the genome databases showed that the aquatic *Phytophthora* species are in fact missing and are thus likely classified as some other *Phytophthora* species. This highlights the poor state of the current databases regarding the fungal/oomycete organisms and underscore that while shotgun sequencing is largely bias free, we still cannot precisely identify many of the community members.

Another major group that can have strong impacts on other organisms are the Chytrids, which are ubiquitous in aquatic environments (Rasconi et al. 2011) and are regarded as indwellers. In our study the proportions of Chytrids in the total fungal community ranged from 3% to 16.1% and this taxon was consistently present in most of the lakes. Chytrids play important roles in ecosystem functioning and food-web dynamics as their zoospores are highly nutritional food sources for zooplankton and may establish alternative trophic links between primary and secondary production in pelagic ecosystems (Kagami et al. 2007, Rasconi et al. 2011, Agha et al. 2016). Chytrids can also remobilize energy by decomposing organic matter (Gleason et al. 2008) and some are also known to be parasites of aquatic plants and animals (Gleason et al. 2014). The species identified in our data included Catenariaceae of the order Blastocladiales which is a known parasite of the pathogenic oomycete *Phythophtora parasitica* (Daft and Tsao 1984) which was in turn one of the most abundant *Phytophthora* species in our dataset. This points to intricate biotic interactions with the fungal community and a complex role of various fungal species in lakes. These findings also emphasize the need for studying the dynamics of these communities in more detail to understand the overall impacts they may have on the lake ecosystem.

## Conclusions

Our exploratory study of freshwater fungal and oomycete communities demonstrated that shotgun sequencing data is suitable for studying the fungal communities and facilitates the analysis of all the different members of the microbial community simultaneously. However, our attempt to classify the pathogenic *Phytophthora* down to species level clearly illustrate the limitations imposed by insufficient species representation of even the most economically important species in current databases and highlights the need to increase reference genome sequencing efforts for microbial eukaryotes. The analysis of fungal communities across the lake dataset suggested that while the community composition varies between lakes, they are distinct from soil communities and likely less impacted by imported fungi as we saw no difference in diversity across the lakes from the different climatic zones. Our survey further suggests that organic substrate availability could be one influential factor shaping fungal communities. As this carbon availability is tightly connection to features of the surrounding watershed, it can be hypothesized that changes in the areas surrounding lakes, such as clear cuts or increasing surface runoff due to climate change, could have a drastic impact on lake fungal communities. We conclude that fungal communities in lakes contribute significantly to microbial diversity in lakes and call for more efforts to increase the coverage of fungal genomic diversity in databases.

## Acknowledgements

We are grateful to the Science for Life Laboratory for funding. The computations were performed on resources provided by the Swedish National Infrastructure for Computing (SNIC), partially funded by the Swedish Research Council through grant agreement no. 2018-05973, through Uppsala Multidisciplinary Center for Advanced Computational Science (UPPMAX) under Project SNIC 2020-5-196. Shotgun sequencing was performed at the Swedish National Genomics Infrastructure (NGI) at Science for Life Laboratory (SciLifeLab) in Uppsala.

---

**Supplementary Figure S1.**
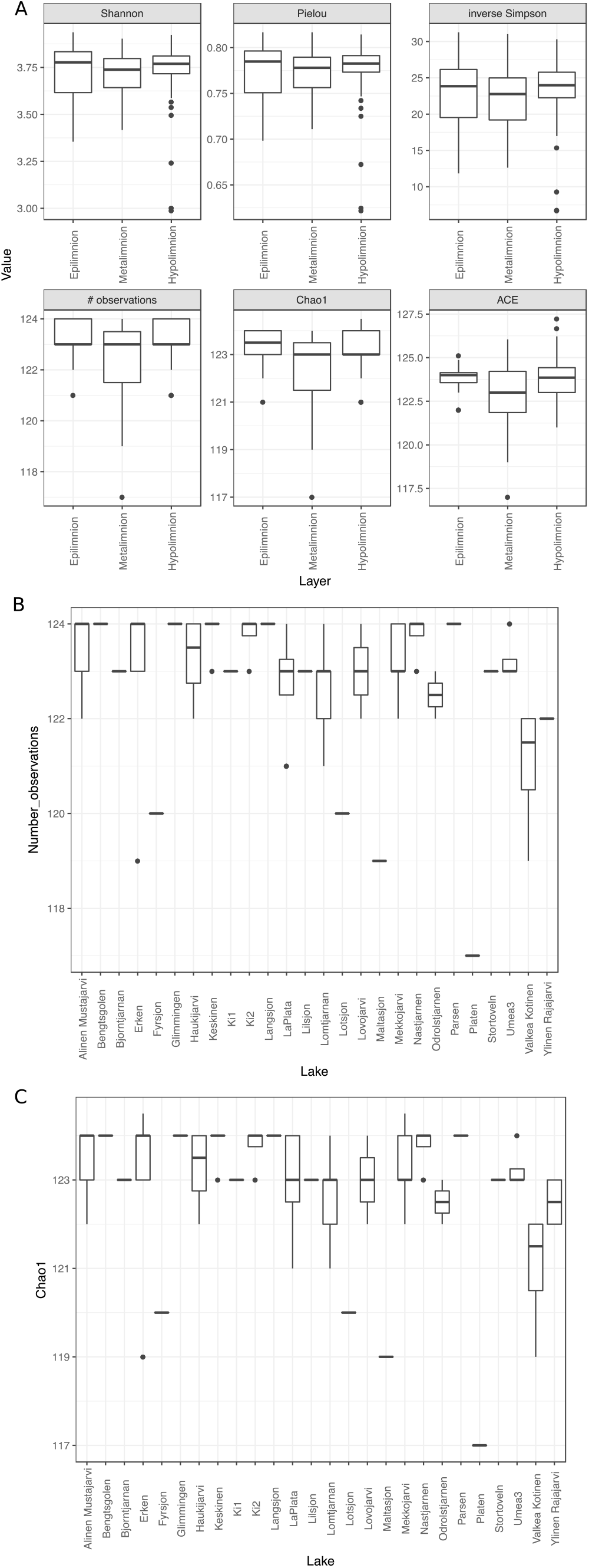
Alpha diversity across: (A) different layers of the lakes, (B) number of observations among lakes, (C) Chao 1 among lakes.

**Supplementary Figure S2.**
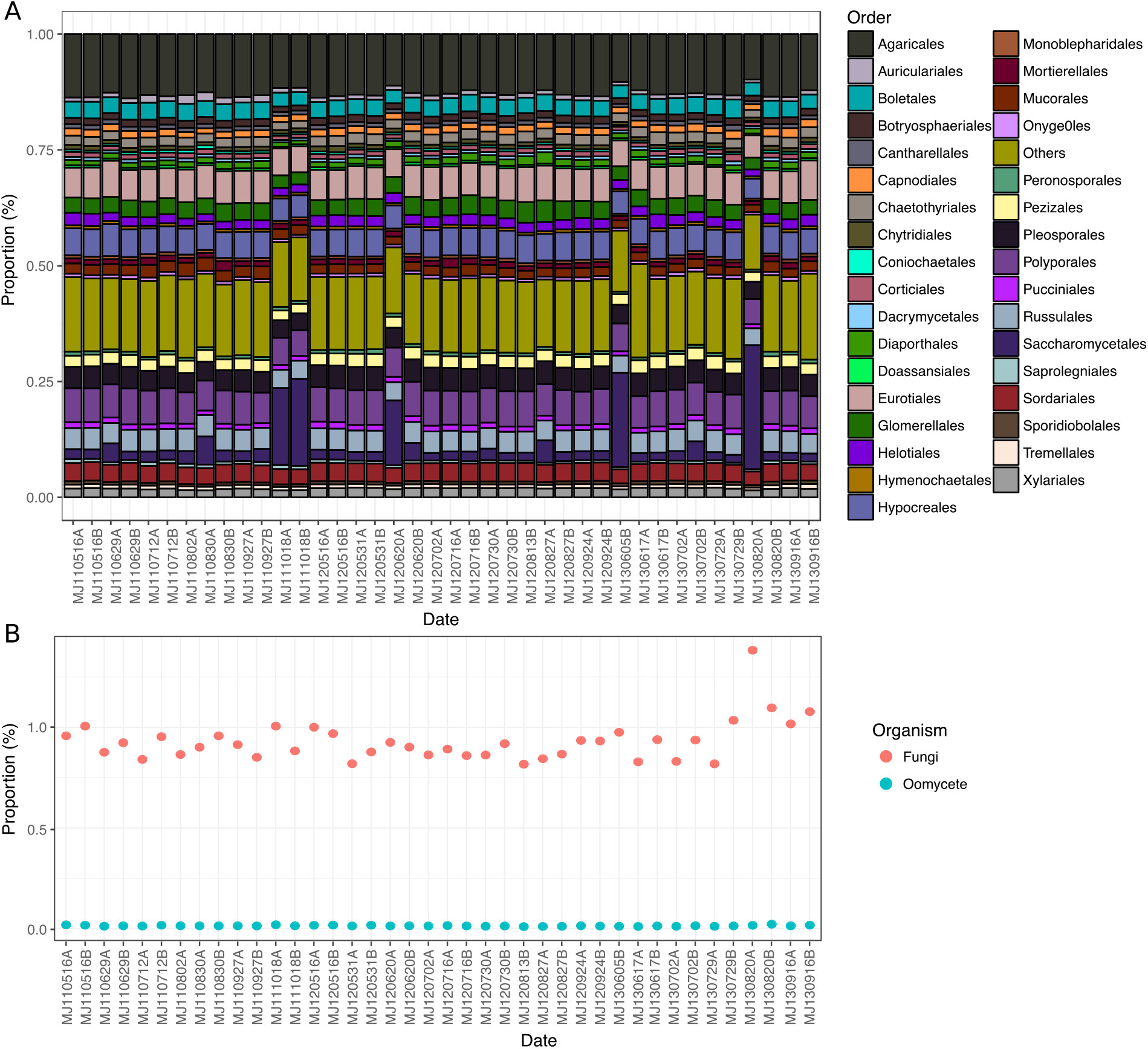
A) The composition of fungal and oomycetes community at order level and B) the proportion of fungal and oomycete reads of all classified reads across time at Lake Mekkojärvi during open water seasons 2011-2013. A and B in the sample names indicate samples taken at the same time point from different layers of the lake.

